# A Genetic and Immunohistochemical Analysis of *H. Pylori* Phenotypes and p27 expression in Adenocarcinoma Patients in Jordan

**DOI:** 10.1101/2022.12.18.520940

**Authors:** Suhaila A. Al-Sheboul, Ahmad Abdul-Razzak Mohammad, Yasemin Shboul, Brent Brown, Ismail I. Matalka

## Abstract

Stomach (gastric) cancer is one of the most prevalent and deadly cancers worldwide and the vast majority of gastric cancers are adenocarcinomas. Based on prior research, there is an association between *Helicobacter pylori (H. pylori)* infection and the frequency of duodenal ulcer, distal gastric adenocarcinoma, mucosa-associated lymphoid tissue lymphoma, and antral gastritis. *H. pylori* virulence and toxicity factors have been identified to significantly influence the clinical outcomes of *H. pylori* infection and gastric adenocarcinoma. However, it is unclear exactly how different strains of *H. pylori* infection affect gastric adenocarcinoma. Current research suggests this involves tumor suppressor genes, like p27, but also *H. Pylori* toxic proteins. Therefore, we quantified known *H. Pylori* genotypes within adenocarcinoma patients to establish the prevalence of known toxins that include cytotoxin-associated gene A (cagA) and vacuolating cytotoxin A (vacA) within patients of variable diagnosis. This analysis used gastrectomy samples which were validated for DNA viability. The incidence of *H. Pylori* in adenocarcinoma patients in Jordan was established to be 54.5% positive (ureA gene) with cagA genotype occurrence at 57.1% but also vacA gene ratios 24.7%:22.1%: 14.3%:14.3%. (vacAs1:vacAs2: vacAm1:vacAm2). We found statistically significant evidence that p27 was dysregulated and suppressed within nearly all *H. Pylori* vacA genotypes but also that 24.6% of *H. Pylori* samples analyzed had a different bacterial genotype and curiously that p27 protein expression was retained in 12% of tested adenocarcinoma *H. Pylori* samples. This is suggestive that p27 could be used as a prognostic indicator but also that an unknown genotype could be contributing to the regulatory effects of p27 protein within this bacterial and cellular environment.

**Key Contribution:** **In Jordan**, this is the first retrospective study to use validated viable gastrectomy tissue samples rather than tissue biopsies to characterize each of the currently defined *H. pylori* strains in patients with variable gastric cancer diagnosis whilst also comparing results with p27 expression within 8 years using these laboratory techniques.

**Conclusions:** Based on these results, a reduction in p27 expression in *H. pylori*-infected men and women of all ages may be associated with the occurrence of gastric adenocarcinomas. Therefore, p27 may represent a key diagnostic marker and predictor of adenocarcinoma prognosis in such individuals and may find use for guiding subsequent treatment decisions.

## 1. Introduction

Stomach cancer (also known as gastric cancer) is the third most common cancer-related cause of death worldwide and 95% are adenocarcinomas [1]. Adenocarcinomas develop from glands of stomach mucosa, or its most superficial layer [1]. However, there are additional cancers that can develop from the stomach, such as leiomyosarcomas, which develop from the muscles that surround the mucosa, and mucosa-associated lymphoid tissue lymphomas, which emerge from the stomach’s lymphoid tissue [1]. Gastric cancers are frequently discovered at advanced stages so the prognoses are often poor [ref}. *H. pylori* was classified as a class 1 carcinogen by the World Health Organisation (WHO) in 1994, with epidemiological, clinical, and experimental data demonstrating a link between *H. pylori* infection and the progression of gastric adenocarcinoma [2]. Patients with *H. pylori* infection have a 6-fold higher chance of developing gastric cancer [3]. The presence of this bacterium increases the risk of developing gastric adenocarcinoma [4]. *H. pylori* possesses virulence factors that affect toxicity and pathogenicity including cytotoxin–associated pathogenicity island (cagPAI) and vacuolating cytotoxin (vacA). VacA enhances *H. Pylori* colonization of the gastrointestinal tract as a pore-forming toxin. In addition, cagPAI contains genes that encode cagA (molecular mass 120–140 kDa), which affects components of the intestinal type IV secretion system, through which cagA protein is delivered into host cells upon bacterial attachment [5]. These toxic factors have been proven to be crucial in defining the clinical outcomes of *H. pylori* infection and the development of gastric adenocarcinoma [5]. Prior laboratory studies have demonstrated the cagA role in carcinogenesis through stimulation of aberrant cell proliferation [6]. Over 90% of *H. pylori* strains occur in southeast Asia with more than 60% of strains from Western populations (including Europe and North America) reported to possess the CagA gene [7]. The VacA gene produces a toxin (p33 / p55 subunits) known to bind to integrin receptors that can interfere with dendritic cells, macrophages and suppress T cells with resulting differential effects on not only regulatory but also effector T cell differentiation; these are essential in immune systems of clinically diagnosed cancer patients and may therefore contribute to prolonged microbial infection thereby affecting disease outcomes [8]. VacA affects this through the nuclear activation of NF-kB and upregulation of an inflammatory cytokine, interleukin-8 (IL-8). The presence of various genotypes of the vacA gene has been reported, including s1a, s1b, s2, ml, and m2 strains of *H. pylori* and they display unusually variable toxicity [9]. In general, strains of type s1a and m1 produce the most toxin, followed by type s1b and type m2 strains, which generate toxin in moderate amounts, while type s2 and type m2 strains exhibit little to no vacuolar toxin activity [10,11]. Therefore, characterizing the toxicity of *H. Pylori* in specific carcinomas allows further knowledge of toxicity factors within clinically diverse and vulnerable patients.

In several prior studies conducted around the world, cagA, vacA s1, and vacA m1 genes, have been thought to play a role in pathogenicity and are linked to gastric cancer [12–14]. The geographic distribution of *H. pylori* strains varies; for instance, the most common strains in East Asia carry the cagA, vacAs1, and vacAm1 genes [15]. The prevalence of the vacAs1b subtype was approximately 100% in South America, 80% in the Spain and Portugal strains, and very low in East Asia [15]. The mechanism of the development of gastric adenocarcinoma in *H. pylori*-infected samples has not yet been determined [16]. One proposed mechanism for the development of cancer is the inactivation of the p27^KIP1^ tumor suppressor gene, located on chromosome 12p13 [17,18]. Eukaryotic cells have a network of regulatory proteins that affect cell cycle control, which precisely regulate and control the progression of the cell through the cell cycle [17]. The cell cycle has checkpoints that are crucial components of the control system that prevent malignancies. Activities of the cell cycle checkpoints are governed by cyclin-dependent kinases (CDKs), a family of protein kinases that bind to several types of regulator proteins known as cyclins [19]. The p27 protein (often referred to as KIP^1^) is a member of the CIP/KIP family of tumor-suppressor proteins [20]. It is a CDK inhibitor (CDKi) that uses biochemical processes to prevent the progression of the cell cycle. Inhibition of CDKs is the mechanism by which p27 inhibits the cell cycle. Cell division, proliferation, and apoptosis are other roles of p27 [21]. In fact, reduced p27 expression has been shown to be a marker of aggressive disease and poor prognosis in several cancers, including colon, breast, malignant melanoma, liver, stomach, lung, and brain tumors [22]. Furthermore, *H. pylori* infection in gastric cancer patients has been associated with reduced p27 expression [23]. In this investigation, we sought to identify the relationship between *H. pylori* infection and p27 expression in gastric cancer tissues in Jordanian patients. We utilized gastrectomy samples rather than tissue biopsies taken from patients with *H. pylori*. These samples were clinically diagnosed with variable adenocarcinoma grades from hospitals in Jordan. *H. Pylori* genotypes of each clinical sample were characterized alongside p27 protein expression confirmation. To ascertain whether p27 protein was related to the development of gastric cancer in these patients, the level of p27 gene expression was also assessed. To our knowledge, this is the only study that has quantified variable *H. Pylori* genotypes in adenocarcinoma with confirmation of expression of p27 protein using samples obtained from patients in Jordan.

## 2. Materials and Methods

### 2.1. Sample Collection

In this study, a total of 77 archived histological samples with linked patient demographic and clinical data were obtained within 8 years (between 2005 and 2013) from the pathology department medical records of King Abdullah University Hospital (KAUH) at Jordan University of Science and Technology (Irbid, Jordan) and the Jordanian Royal Medical Services (Amman, Jordan) (Table 1). These clinical samples (*n* = 77) were examined using fixed paraffin-embedded tissue analysis. All samples were classified as gastric adenocarcinomas and were stored at 25 °C. This study was carried out with consent from the Institutional Review Board (IRB), Ethics Committee, at Jordan University of Science and Technology.

### 2.2. Clinical Samples Processing

Using a microtome, all formalin-fixed tumor specimens that were embedded in paraffin blocks were sectioned into 5-micrometer thick tissue slices (Energy beam science, East Granby, CT, USA). The microtome blade was changed between samples, and its cutting surface was cleaned and sterilized with xylene and absolute ethanol, respectively. 5–10 tissue sections were collected in 1.5 ml sterile Eppendorf nuclease free tubes for DNA extraction, and the 5 μm tissue slices were mounted on positively charged slides for immunohistochemistry.

### 2.3. Immunohistochemistry (IHC) Staining

To evaluate *p27^KIP1^* expression in tissue (s), all tissue section slides were treated with a mouse monoclonal Anti-Human IgG *p27^KIP1^* (Clone SX53G8.5 dilution 1:100; Code *M7203;* Dako Cytomation, Denmark) that cross-reacts with p27^kip1^ using a protocol described previously [24]. Slides were deparaffinized with xylene twice, for 5 min each, rehydrated through a series of graded alcohol washes (100% alcohol, 2 times, for 3 min each, and then transferred once through 95%, 70%, and 50% alcohols for 3 min each). Endogenous peroxidase activity was blocked by incubating sections in 3% H2O2 solution in methanol at room temperature for 10 min. Then slides were rinsed with PBS twice, for 5 min each. To reveal the antigenic epitope, we performed antigen retrieval by pouring 300 mL of 10 mM citrate buffer, pH 6.0 into the staining container containing arranged slides, and incubating it at 95 °C for 23 min using a pt link pretreatment system (Dako, Agilent, Glostrup, Denmark). We removed the staining container at room temperature and allowed the slides to cool for 20 min. This was followed by washing in Dulbecco’s phosphate-buffered saline (PBS) twice, for 5 min each (Sigma Aldrich, St. Louis, MO, USA). Then, we drained the blocking buffer from the slides. All the slides were treated with p27 monoclonal antibody (H-1): sx-53G8.5 (Dako Cytomation, Agilent), diluted 1:100 in Dako antibody diluent, for 45 min at room temperature. We washed the slides with PBS twice, for 5 min each, according to the manufacturer’s recommendations [24]. The detection was carried out using Dako EnVision^®^+ Dual Link System-HRP (DAB+) (Dako, Agilent). The Liquid DAB+ Substrate Chromogen System (Dako, Agilent) was used to view the slides, and Mayer’s hematoxylin (Polysciences) was used as a counterstain. As a negative control, primary antibodies were excluded. Tonsil and lymphoma patient tissue was used as a positive control. More than 5% of neoplastic cells must exhibit prominent brown nuclear staining for samples to be deemed positive for p27 protein expression. The scoring system used to evaluate the results of immunostaining was described previously [25]. To verify the initial diagnosis, all samples were examined and confirmed by a licensed pathologist at the pathology division of KAUH. All cells were counted on the slides using a high-power field scanner, and pictures were taken with a digital camera (Olympus model C-5060, Olympus, Tokyo, Japan).

### 2.4. DNA Extraction

For extracting genomic DNA, 5–10 sections from each block were taken and put into sterile, nuclease-free Eppendorf tubes. Prior to genomic DNA extraction, xylene and paraffin residues from paraffin sections were removed in the pathology lab in accordance with standard operating procedures [26]. To quickly and effectively deparaffinize the tissue, 1 mL of xylene was added to each tube, and the tubes were vortexed violently for 10 seconds. The mixture was then centrifuged at a full speed of 4000 rpm for 2 min at room temperature, with the pellet remaining in the tube. This procedure was performed twice. One milliliter of 96% ethanol was added to each pellet, it was vortexed and centrifuged at full speed for 2 min at room temperature, and the supernatant was removed by pipetting. To remove any leftover xylene, this step was repeated. Next, Eppendorf tubes containing the tissue pellets were opened and incubated at 37 °C for 15–30 min until the ethanol evaporated. Each pellet was then subjected to genomic DNA extraction using a QIAamp DNA FFPE Tissue kit (Qiagen, Hilden, Germany) according to the manufacturer’s instructions. To ensure DNA purity, the extracted DNA was then eluted, and its concentration assessed using a NanoDrop ND-1000 Spectrophotometer (Thermo Scientific, Waltham, MA, USA). For later use, all the DNA extracts were kept in storage at −20 °C. Using an antibody to Ki-67, adjacent sections were immunostained to detect proliferating cells, and apoptotic cells were detected using a terminal deoxynucleotidyltransferase-mediated dUTP-biotin nick end labeling assay. Proliferation and apoptotic indices were then calculated as previously described [24].

### 2.5. Detection of H. pylori Virulence Genes

*H. pylori* virulence genes were detected by PCR amplification. The primers used in this study were described previously (Table 2). The primers targeted the amplification of *H. pylori* genes and the glyceraldehyde 3-phosphate dehydrogenase (GAPDH) gene to serve as an indicator of DNA extraction validity [27].

**Table 1.**
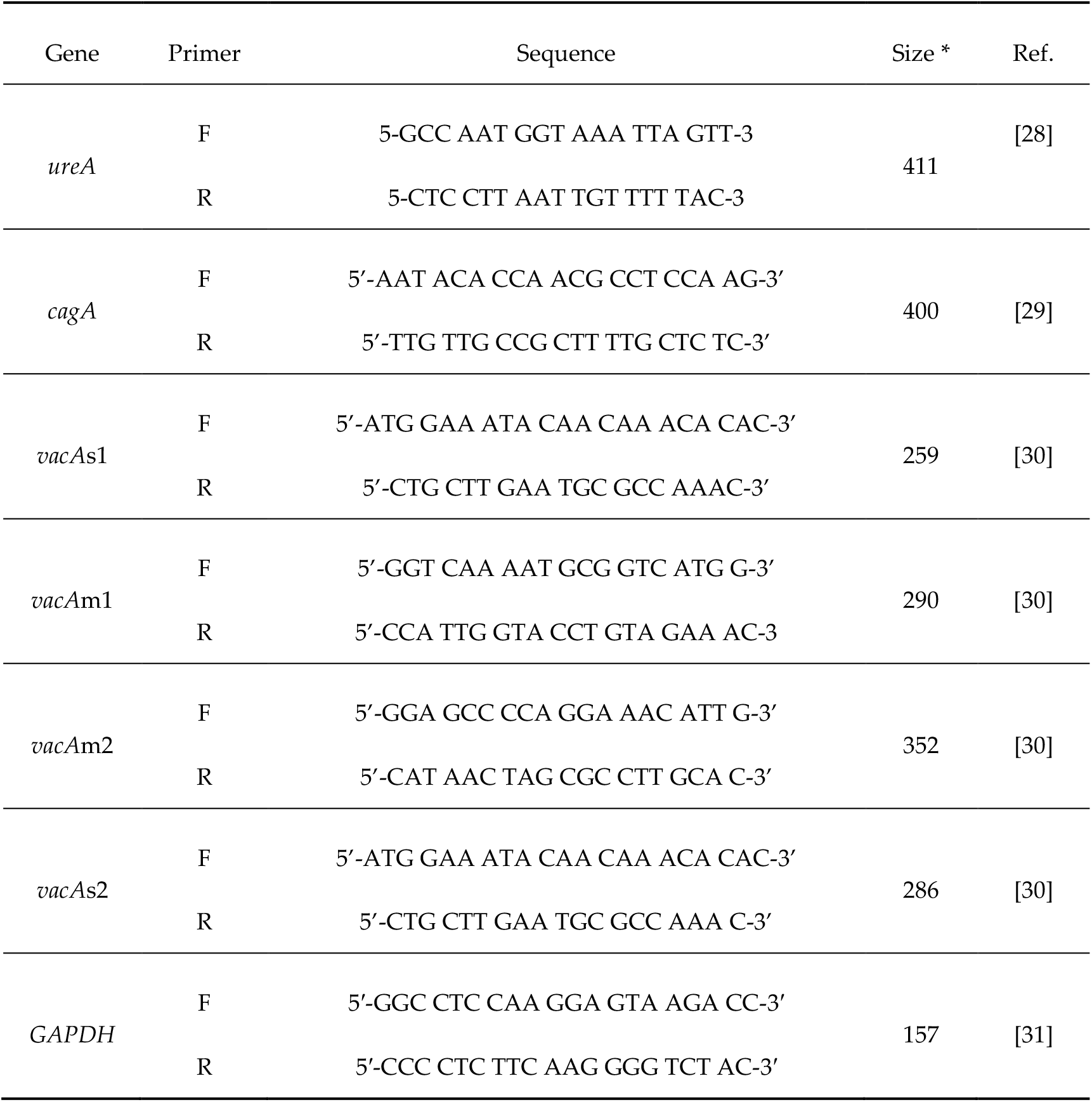
PCR primers for *H. pylori* genes (and GADPH quality control bottom)

### 2.6. PCR for the Glyceraldehyde 3-Phosphate Dehydrogenase (GAPDH) Gene

One of the universal housekeeping genes used to assess the integrity of DNA samples is GAPDH as an internal control. This gene was subjected to PCR. Each PCR reaction with a total volume of 25 μL contained 12.5 μL of PCR master mix (Promega; Madison, WI, USA), 8.0 μL of nuclease free water, 1.0 μL (5 pmol/μL for GAPDH) of each primer, and 2.5 μL of DNA (100 ng/μL). A negative control reaction (containing all components except DNA template) was included. The amplification protocol was run with the thermal profile recommended by the master mix manufacturer; initial denaturation at 95 °C for 5 min, followed by 40 cycles of denaturation at 95 °C for 1 min, annealing at 54 °C for 1 min, and extension at 72 °C for 1 min. The amplification ended with a final extension at 72 °C for 5 min.

### 2.7. PCR ureA Gene to Confirm H. pylori Identity

PCR for the detection of the *H. pylori* ureA gene was performed in 25 μL volumes as described above, using specific *H. pylori* ureA primers along with positive and negative controls. The amplification protocol was run with the thermal profile recommended by the manufacturer; initial denaturation at 95 °C for 10 min, followed by 35 cycles of denaturation at 94 °C for 1 min, annealing at 47 °C for 1 min, and extension at 72 °C for 1 min.

### 2.8. PCR for H. pylori Virulence Genes

Samples that were positive for the ureA gene underwent five PCR cycles to detect the presence of the virulence genes of *H. pylori* (cagA, vacAs1, vacAs2, vacAm1, and vacAm2). PCR reactions were performed in 25-μL volumes. Table 2 displays the temperature profile for each gene PCR cycle.

**Table 2.**
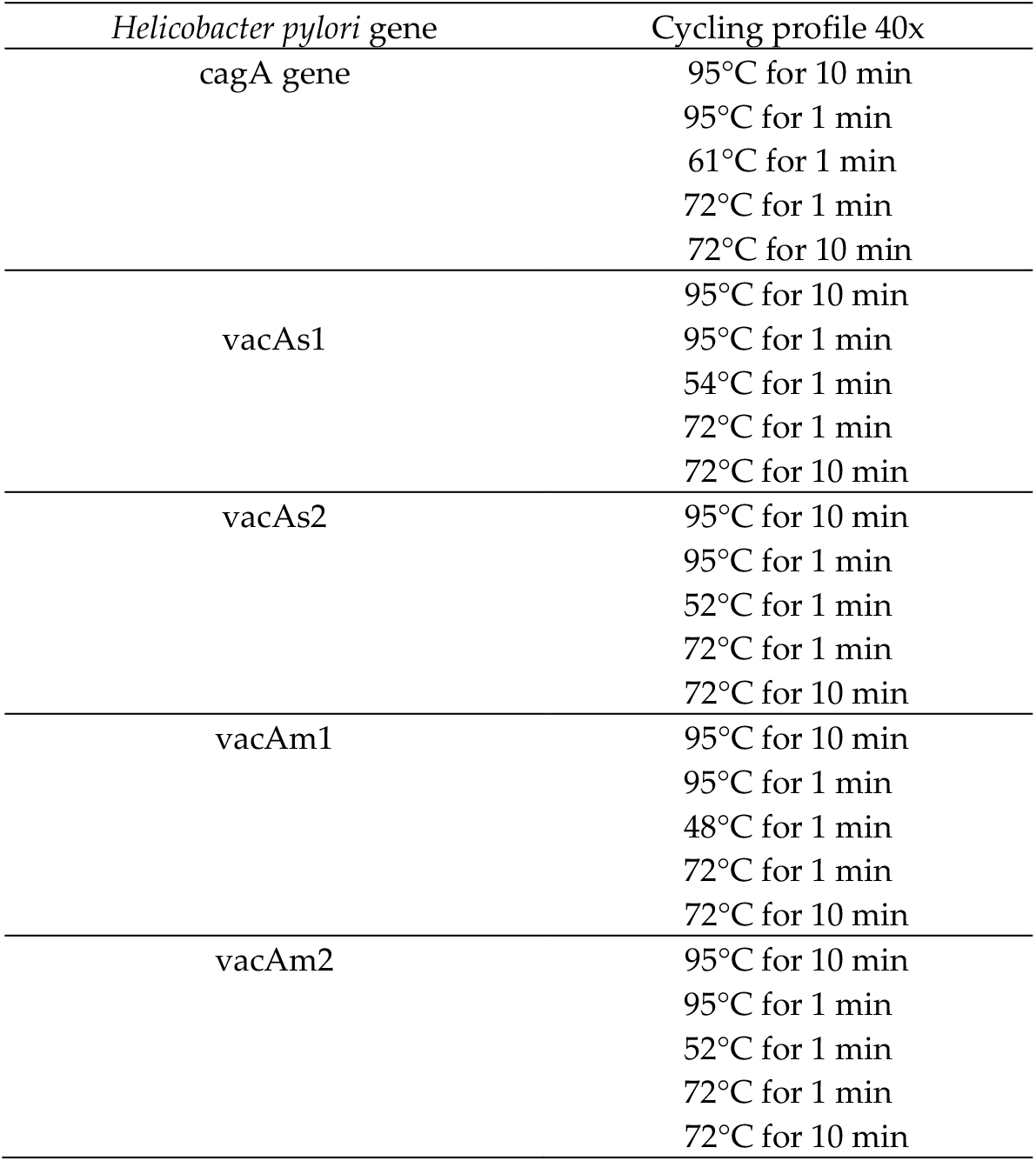
PCR protocol for *Helicobacter pylori* virulence gene amplification.

| *Helicobacter pylori* gene | Cycling profile 40x |
| --- | --- |
| *cagA* gene | 95°C for 10 min |
|  | 95°C for 1 min |
|  | 61°C for 1 min |
|  | 72°C for 1 min |
|  | 72°C for 10 min |
| *vacAs1* | 95°C for 10 min |
|  | 95°C for 1 min |
|  | 54°C for 1 min |
|  | 72°C for 1 min |
|  | 72°C for 10 min |
| *vacAs2* | 95°C for 10 min |
|  | 95°C for 1 min |
|  | 52°C for 1 min |
|  | 72°C for 1 min |
|  | 72°C for 10 min |
| *vacAm1* | 95°C for 10 min |
|  | 95°C for 1 min |
|  | 48°C for 1 min |
|  | 72°C for 1 min |
|  | 72°C for 10 min |
| *vacAm2* | 95°C for 10 min |
|  | 95°C for 1 min |
|  | 52°C for 1 min |
|  | 72°C for 1 min |
|  | 72°C for 10 min |

The resulting amplicons and their corresponding lengths were visualized and documented via gel electrophoresis and analysis (Quantity 1 software, Bio-rad, CA, USA).

### 2.9. Statistical Analysis

All data were analyzed using SPSS version 19.0 software (SPSS, Inc., Chicago, IL, USA). Pearson’s chi-square or Fisher’s exact tests were used to analyze the relationship between *H. pylori* infection, virulence genes, and p27 gene expression in the gastric cancer patients analyzed. At *p* ≤ 0.05, the results were considered significant.

### 2.10. Ethical Statement

This study was approved by the University Review Committee for Research on Humans of the King Abdullah University Hospital, and the University Research Committee of the Jordan University of Science and Technology

## 3. Results

### 3.1. Demographic and Clinical Data

Patient ages ranged 30 to 98 years at diagnosis; 35 were females, and 42 were males. Of the 77 gastric cancer cases, 39 were differentially diagnosed with diffuse type cancer and 38 were diagnosed with intestinal type. Table 3 summarizes the demographic and clinical information for the two patient groups.

**Table 3.** Gastric sample classification.

| Sample | Year | Age | Sex | Diagnosis | Type |
| --- | --- | --- | --- | --- | --- |
| 1 | 2003 | 59 | Female | moderately differentiated adenocarcinoma | Diffuse |
| 2 | 2003 | 61 | Male | moderate chronic gastritis. | Intestinal |
| 3 | 2003 | 59 | Male | invasive moderately differentiated adenocarcinoma | Diffuse |
| 4 | 2004 | 76 | Male | moderately differentiated adenocarcinoma | Intestinal |
| 5 | 2004 | 54 | Male | invasive adenocarcinoma | Diffuse |
| 6 | 2004 | 83 | Male | invasive moderately to poorly differentiated adenocarcinoma | Diffuse |
| 7 | 2004 | 84 | Male | moderately differentiated adenocarcinoma | Diffuse |
| 8 | 2004 | 43 | Male | poorly differentiated adenocarcinoma | Diffuse |
| 9 | 2005 | 50 | Female | invasive moderately differentiated adenocarcinoma | Intestinal |
| 10 | 2005 | 81 | Male | moderately differentiated adenocarcinoma | Diffuse |
| 11 | 2005 | 66 | Female | poorly differentiated adenocarcinoma | Diffuse |
| 12 | 2005 | 33 | Female | invasive differentiated adenocarcinoma | Intestinal |
| 13 | 2005 | 45 | Female | poorly differentiated invasive adenocarcinoma | Diffuse |
| 14 | 2005 | 71 | Male | invasive moderately differentiated adenocarcinoma | Intestinal |
| 15 | 2005 | 51 | Female | invasive poorly differentiated adenocarcinoma | Diffuse |
| 16 | 2005 | 44 | Female | invasive differentiated adenocarcinoma | Intestinal |
| 17 | 2006 | 98 | Female | poorly differentiated adenocarcinoma | Intestinal |
| 18 | 2006 | 51 | Female | poorly differentiated adenocarcinoma | Diffuse |
| 19 | 2006 | 83 | Male | poorly differentiated Invasive gastric adenocarcinoma | Intestinal |
| 20 | 2007 | 54 | Male | poorly differentiated adenocarcinoma | Diffuse |
| 21 | 2007 | 69 | Female | poorly differentiated adenocarcinoma | Intestinal |
| 22 | 2007 | 79 | Male | poorly differentiated adenocarcinoma | Diffuse |
| 23 | 2007 | 52 | Female | poorly differentiated adenocarcinoma with ulcerated surface | Intestinal |
| 24 | 2007 | 73 | Male | moderately differentiated adenocarcinoma | Diffuse |
| 25 | 2007 | 76 | Female | moderately differentiated adenocarcinoma | Intestinal |
| 26 | 2007 | 82 | Male | poorly differentiated adenocarcinoma | Intestinal |
| 27 | 2007 | 75 | Male | poorly differentiated invasive adenocarcinoma | Diffuse |
| 28 | 2007 | 47 | Male | poorly differentiated adenocarcinoma with large area of necrosis | Diffuse |
| 29 | 2007 | 67 | Female | poorly differentiated adenocarcinoma | Diffuse |
| 30 | 2007 | 77 | Male | moderately differentiated adenocarcinoma | Intestinal |
| 31 | 2008 | 37 | Female | poorly differentiated adenocarcinoma | Diffuse |
| 32 | 2008 | 52 | Female | moderately differentiated adenocarcinoma | Intestinal |
| 33 | 2008 | 71 | Male | moderately differentiated adenocarcinoma | Intestinal |
| 34 | 2008 | 30 | Female | poorly differentiated gastric adenocarcinoma | Diffuse |
| 35 | 2008 | 42 | Female | poorly differentiated adenocarcinoma | Diffuse |
| 36 | 2009 | 44 | Female | poorly differentiated adenocarcinoma with endocrine differentiation | Diffuse |
| 37 | 2009 | 48 | Male | poorly differentiated adenocarcinoma | Diffuse |
| 38 | 2009 | 52 | Male | moderately differentiated ulcerated adenocarcinoma | Intestinal |
| 39 | 2009 | 58 | Female | poorly differentiated adenocarcinoma | Intestinal |
| 40 | 2009 | 75 | Male | moderately differentiated adenocarcinoma | Intestinal |
| 41 | 2009 | 46 | Male | moderately differentiated adenocarcinoma | Intestinal |
| 42 | 2009 | 77 | Female | poorly differentiated gastric adenocarcinoma | Diffuse |
| 43 | 2010 | 80 | Female | poorly differentiated adenocarcinoma reaching subserosa | Intestinal |
| 44 | 2010 | 55 | Female | poorly differentiated adenocarcinoma | Intestinal |
| 45 | 2010 | 43 | Female | poorly differentiated adenocarcinoma | Diffuse |
| 46 | 2010 | 50 | Female | moderately differentiated adenocarcinoma | Intestinal |
| 47 | 2010 | 49 | Female | poorly differentiated adenocarcinoma | Diffuse |
| 48 | 2010 | 56 | male | poorly differentiated adenocarcinoma | Intestinal |
| 49 | 2011 | 75 | male | moderately differentiated adenocarcinoma | Intestinal |
| 50 | 2011 | 67 | Female | poorly differentiated adenocarcinoma | Diffuse |
| 51 | 2011 | 40 | male | poorly differentiated adenocarcinoma | Intestinal |
| 52 | 2012 | 61 | male | poorly differentiated adenocarcinoma | Intestinal |
| 53 | 2012 | 32 | Female | poorly differentiated adenocarcinoma | Diffuse |
| 54 | 2012 | 62 | Female | poorly differentiated adenocarcinoma | Diffuse |
| 55 | 2010 | 68 | male | poorly differentiated gastric adenocarcinoma | Diffuse |
| 56 | 2009 | 40 | Female | poorly differentiated adenocarcinoma | Diffuse |
| 57 | 2009 | 70 | Female | poorly differentiated adenocarcinoma | Diffuse |
| 58 | 2012 | 81 | male | poorly differentiated gastric adenocarcinoma | Diffuse |
| 59 | 2008 | 78 | male | moderately differentiated adenocarcinoma | Intestinal |
| 60 | 2010 | 72 | male | moderately differentiated adenocarcinoma | Intestinal |
| 61 | 2010 | 45 | Female | poorly differentiated adenocarcinoma | Diffuse |
| 62 | 2011 | 70 | Female | moderately to poorly differentiated adenocarcinoma | Intestinal |
| 63 | 2012 | 69 | male | poorly differentiated adenocarcinoma | Diffuse |
| 64 | 2012 | 86 | male | moderately differentiated adenocarcinoma | Intestinal |
| 65 | 2011 | 68 | Female | moderately to poorly differentiated adenocarcinoma | Intestinal |
| 66 | 2010 | 70 | male | moderately differentiated adenocarcinoma | Intestinal |
| 67 | 2012 | 66 | male | poorly differentiated adenocarcinoma | Diffuse |
| 68 | 2010 | 52 | male | moderately differentiated adenocarcinoma | Intestinal |
| 69 | 2010 | 65 | Female | poorly differentiated adenocarcinoma | Intestinal |
| 70 | 2012 | 63 | male | moderately differentiated adenocarcinoma | Intestinal |
| 71 | 2010 | 51 | male | poorly differentiated adenocarcinoma | Diffuse |
| 72 | 2012 | 58 | male | poorly differentiated adenocarcinoma | Diffuse |
| 73 | 2012 | 83 | male | poorly differentiated adenocarcinoma | Diffuse |
| 74 | 2010 | 60 | male | poorly differentiated adenocarcinoma | Intestinal |
| 75 | 2011 | 50 | Female | poorly differentiated adenocarcinoma | Diffuse |
| 76 | 2013 | 75 | male | moderately differentiated adenocarcinoma | Intestinal |
| 77 | 2012 | 54 | male | poorly differentiated adenocarcinoma | Diffuse |

### 3.2. Immunohistochemistry Results

Demonstrations of the positive immunohistochemistry control, and negative and positive staining performed using antibodies against p27^kip1^ low power (10X), are presented in Figures 1–3. Immunohistochemistry indicated that 56/77 (72.7%) of the gastric cancer samples were negative for *p27* gene expression. Of the 56 negative samples, 27 were from females and 29 from males. Thirty cases of the negative IHC samples were of diffuse-type gastric cancer, while the remaining 26 were intestinal type. The cellular staining between *p27* expression and *H. pylori* genes is shown below.

**Figure 1.**
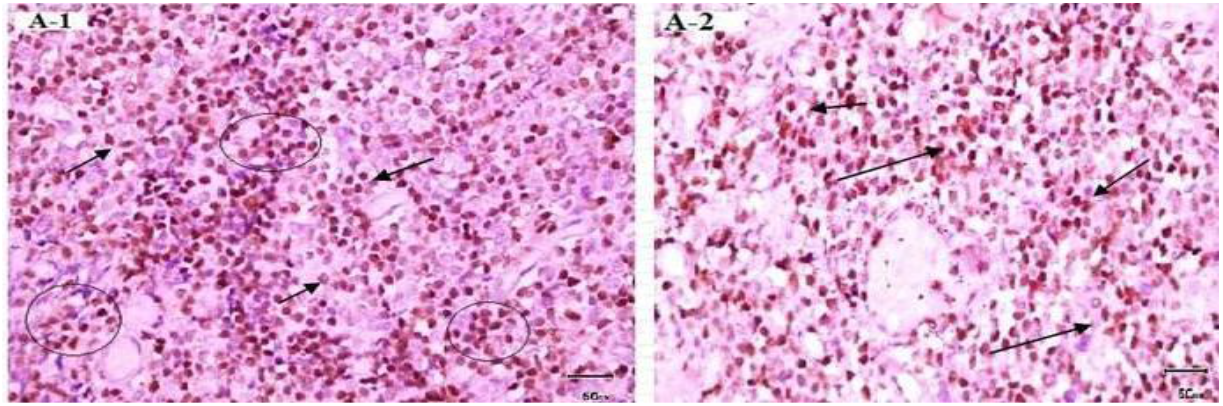
Immunohistochemistry staining of positive controls for p27. A-1: Positive control from tonsil tissue section showing cell staining (brown) of tonsil tissue sample (10x). A-2: Positive control from lymphoma tissue showing cell nuclei staining (brown) in lymphoma sample (40x). Primary antibody anti-p27, secondary antibody anti-Ki67 with peroxidase, DAB chromogen and hematoxylin stain.

**Figure 2.**
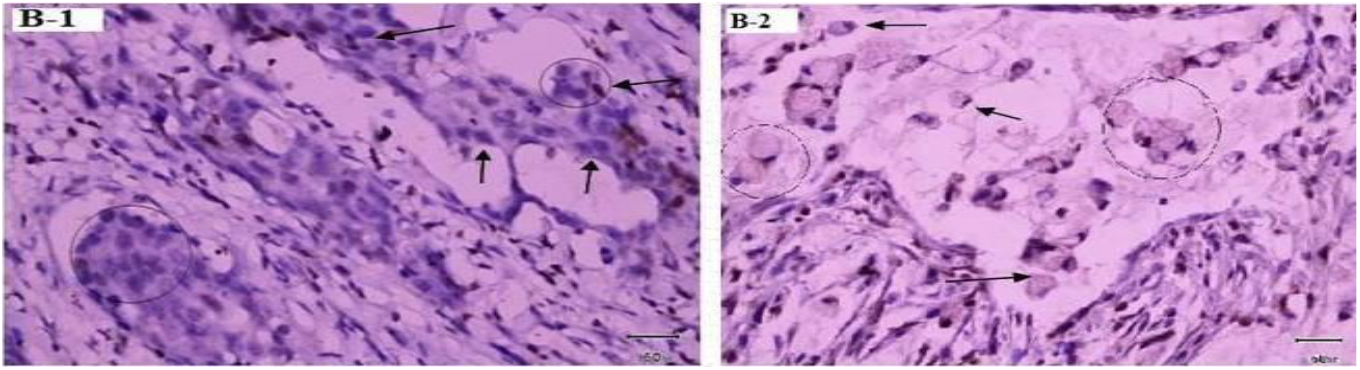
Immunohistochemistry of negative control for P27 expression in cancer tissue. B-1: Negative control from H. Pylori uninfected gastric intestinal adenocarcinoma sample showing nuclei staining only. Low power (10x) view. B-2: Negative control from H. Pylori un-infected gastric diffuse adenocarcinoma showing blue nuclear stain around gastric diffuse adenocarcinoma sample. Low power (10x) view. No primary antibody, secondary antibody anti-Ki67 with peroxidase, DAB chromogen and hematoxylin stain

**Figure 3.**
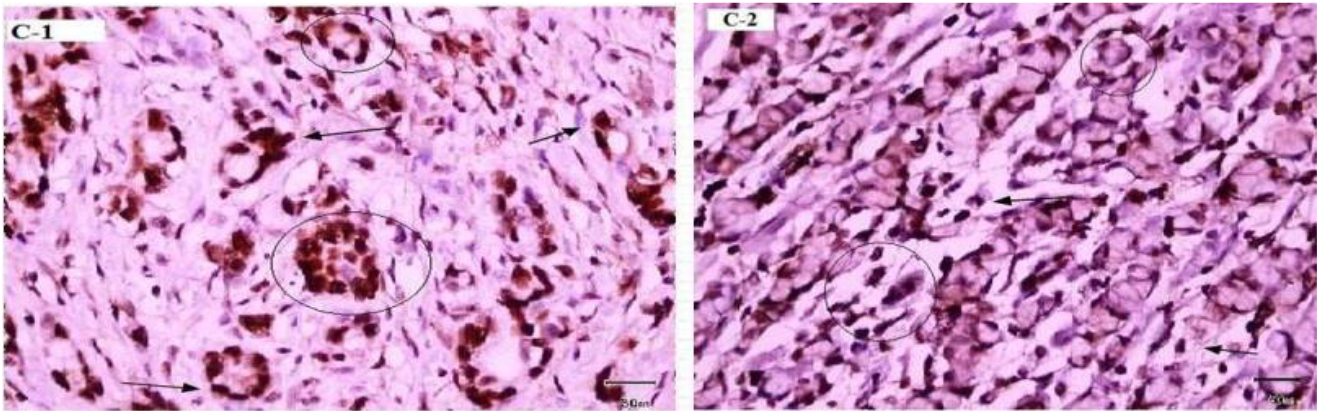
Immunohistochemical Expression of p27 and Ki67 in Cancer Cells. C-1: Immunohistochemical detection of p27 and Ki67 expression. Low power (10x) view of clear positive (brown stain) in epithelium of H. Pylori infected gastric intestinal adenocarcinoma sample. C-2: Immunohistochemical detection of p27kip1p27 and Ki67 expression. Low power (10x) view of clear positive (brown stain) in epithelium of H. Pylori infected of gastric diffuse adenocarcinoma sample. Primary antibody anti-p27, secondary antibody anti-Ki67 with peroxidase, DAB chromogen and hematoxylin stain.

### 3.3. PCR Results

All samples were positive for GAPDH, indicating that the test samples contained high-quality DNA and were suitable for further analysis. Regarding the ureA gene, 42 out of 77 samples were positive for the *H. pylori* ureA gene; however, no correlations between the ureA gene presence and the cancer type or gender were found. The majority of samples (37/42) positive for the ureA gene were negative for p27 protein expression (Table 4), with statistical significance (p<0.05) indicative that that the p27 protein was suppressed in almost all gastric cancer samples that were positive for *H. pylori* infection and ureA gene. The presence of virulence genes and their combinations are summarized (Table 4).

**Table 4.** Association between PCR of ureA gene and IHC results of p27 expression.

| P27 expression | H. pylori Detection |  | Total |
| --- | --- | --- | --- |
|  | Negative ureA gene | Positive ureA gene |  |
| IHC Negative for p27 | 19 | 37 | 56 |
| IHC Positive for p27 | 16 | 5 | 21 |
| Total | 35 | 42 | 77 |
| Statistical significance | p value >0.05 | p value <0.05 | p value <0.05 |

**Table 5.** *H. pylori* (ureA+) (vacA+), genotypes in gastrectomy tissue samples (*p*-value > 0.05)

| Genotype | Prevalence (%) |
| --- | --- |
| vacA s1 (n=77) | 19 (24.7%) |
| vacA s2 (n=77) | 17 (22.1%) |
| vacA m1 (n=77) | 11 (14.3%) |
| vacA m2 (n=77) | 11 (14.3%) |
| ureA-vacA- (n=77) | 19 (24.6%) |

**Table 6.** Associations of virulence genes with p27 gene expression (*p*-value < 0.05)

| Positive Virulence Gene | P27 expression |  | P value |
| --- | --- | --- | --- |
|  | Positive | Negative |  |
| cagA (n=24) | 1 | 23 | <0.05 |
| vacAs1 (n=19) | 1 | 18 | <0.05 |
| vacAs2 (n=17) | 3 | 14 | <0.05 |
| vacAm1 (n=11) | 0 | 11 | <0.05 |
| vacAm2 (n=11) | 0 | 11 | <0.05 |

No statistically significant correlations between any gene occurrence, gender, or type of gastric cancer were found in this histological analysis; however all virulence genes demonstrated significance (*p*-value < 0.05) correlation with protein p27 expression based on IHC and PCR data (Table 6).

## 4. Discussion

*H. pylori* is one of the most common bacterial infections in humans that may progress to gastritis, duodenal ulcers, peptic ulcer disease (PUD), gastric adenocarcinoma, and mucosa-associated lymphoid tissue (MALT) lymphoma [2]. In the current study, we investigated the potential correlation between *H. pylori* infection and gastric adenocarcinoma in a Jordanian population. Although several studies have been conducted in Jordan to investigate the relationship between *H. pylori* infection and gastric cancer [32–35], to the best of our knowledge, this is the first study of its kind in the Middle East to look at the prevalence of *H. pylori* in patients with gastric cancer using gastrectomy samples rather than biopsies.

Seventy-seven samples of adenocarcinoma gastrectomy, formalin-fixed, paraffin-embedded tissue samples were collected from the archives of the Pathology Department at Jordanian Royal Medical Services and King Abdullah University Hospital (KAUH) (Amman and Irbid cities, respectively) (Table 1). The samples were examined for the presence of past *H. pylori* infection, the prevalence of cagA and vacA allelic subtypes, and their correlation with one another, as well as with the expression of the tumor suppressor protein, p27.

We confirmed the presence of *H. pylori* in 42 out of 77 (54.5%) gastrectomy samples, using PCR amplification of the ureA gene (Table 5), while the cagA virulence gene was detected in these samples in 57.1% (24/42) of cases. A total of 42 samples positive for the ureA gene were further processed for the amplification of cagA and vacA alleles (s1, s2, m1, m2) (Table 5). According to our study, the predominant vacA genotypes of patients with *H. pylori* infection in the Jordanian population were s1: 24.7%, s2: 22.1%, m1: 14.3%, and m2 positive: 14.3%. On the other hand, we investigated the link between the expression of p27 and *H. pylori* infection. Our results showed that 56/77 (72.7%) of gastric adenocarcinomas tested negative for p27 expression (Table 6) (Figure 3). Furthermore, 37 of 42 gastric adenocarcinoma tissue samples that were positive for ureA were also negative for p27 expression, with a significant correlation (*p*-value 0.05) (Table 4). Studies showed that cagA positivity rates and their clinical outcomes differ from one country to another and among different population groups within the same country [36].

In general, the cagA prevalence rate has been found to be 50–60% in Middle Eastern countries [37,38]. In our study, the cagA prevalence rate was 57.1%, whereas 100% of *H. pylori* strains in East Asian countries are cagA-positive [39]. From our results, we can demonstrate that virulent *H. pylori* strains associated with adenocarcinoma often carry cagA, as more than half of the cases under study were positive for the *H. pylori* strain carrying the cagA gene. Our results agree with those of a study performed in Japan, Korea, the United States, and Colombia that reported predominance of the cagA genotypes [36,37,38–42]. The different combinations of vacA s and m regions identify the virulence characteristics of the *H. pylori* strain. It has been shown that type s1m1 strains produce higher cytotoxin activity *in vitro* than type s1m2 strains, whereas the s2m2 strains do not produce detectable amounts of the cytotoxin and thus are considered less virulent [43]. Therefore, it is significant to identify the vacA profiles of the isolated strains and then evaluate the subtype combinations, together with the clinical outcome of the patient. According to the results of our study, the predominant genotype was found to be vacAm1s1—unlike in other Middle Eastern countries, e.g., the predominant subtype was reported to be vacAm1s2 in Turkey [38]; but similar to the results of published reports from other countries like South Africa [41] and Mexico [44]. The study of genotypes in four different countries reported that the vacAs1ml genotypes were predominant in Japan, Korea, the United States, and Colombia [36]. The same study reported a higher prevalence of the vacA s1 than the vacA s2 genotype. A study from Germany showed the most frequent allelic combinations were s1m2: 47.7%, s1m1: 35.4%), and s2m2: 15.4% [42]. Our results showed that 72.7% of gastric adenocarcinoma samples were negative for p27 expression, which means that the lack of p27 expression can be an essential change during gastric carcinogenesis. More recent research indicates differential resistance of anti-microbial therapeutics worthy of further investigation between cagA positive and vacA positive pylori [45].

Our results confirm a strong correlation between the presence of *H. pylori* in gastric adenocarcinoma and a lack of p27 expression as 37 out of 42 samples with positive ureA were negative for p27 expression with a significant association (p = 0.001). This result is consistent with a study done in vitro by Shirin et al., who found a strong correlation between the presence of *H. pylori* and the inhibition of p27 immunoexpression [46]. Although prior studies have indeed found a correlation between different carcinoma types and expression of p27, we believe this is the first study to document this using validated viable gastrectomy samples. Therefore, this method of ascertaining individual phenotypes is viable to determine historical *H. Pylori* strains. Furthermore, all samples were collected between 2003 – 2013 and validated by PCR and immunohistochemical analysis occurring between 2005 – 2013. Existing protocols at the time of project were indicative that longer PCR primers could potentially be analyzed for up to 20 years and antigenic degradation can occur of immunohistochemical samples similarly (47—50). Whereas at that time we utilized an additional GAPDH assay to quantify DNA quality with guidelines on next generation sequencing. In our combined analysis we show that FFPE could potentially be valid in combination with PCR utilizing FFPE immunohistochemistry on gastrectomy samples between 8 to 10 years. In addition we utilized a well characterized ki67 protein marker IHC as a standard known marker of cell proliferation (51).

## 4. Limitations

PCR testing of unknown phenotypes; testing of gastric adenocarcinoma precursor condition tissues to assess p27 expression and other results; stratified or regression analyses by demographics, cancer sub-type, prior treatment of *H. pylori* with antibiotics.

## 5. Conclusion

We concluded that the suppression of p27 expression and *H. pylori* infection were correlated. Several earlier studies demonstrated a positive correlation, and their results are all consistent with one another. Interestingly 24.6% of H. Pylori genotypes sampled were unknown and 12% retained positive p27 protein expression indicating that it is possible that certain *H. Pylori* unknown phenotypes could play a positive regulatory role in p27 protein expression which requires further study. The expression of the vacA genotype does not however, appear to be associated with the inhibition of p27. The absence of p27 expression was found to have a propensity for the cagA (+) *H. pylori* genotype, despite the negative correlation. Additional research would help to firmly establish this association and pave the way for new perspectives for diagnosis, prognostic biomarkers, and targeted or individualized therapeutics.

## Supplementary Materials

The following supporting information can be downloaded at: Figure S1-14: title; Table S1-S12: Further information on paraffin and formalin embedding; Paraffin Embedding Technique;A Review of Preanalytical Factors Affecting Molecular, Protein, and Morphological Analysis of Formalin-Fixed, Paraffin-Embedded (FFPE) Tissue: How Well Do You Know Your FFPE Specimen? (2014) | Archives of Pathology & Laboratory Medicine (allenpress.com); Immunofluorescence Staining of Paraffin Sections Step by Step (2020).

## Author Contributions

Conceptualization, S.A.-S.; methodology, A.A.-R.M.; software, S.A.-S. and A.A.-R.M.; validation, S.A.-S. and A.A.-R.M.; formal analysis, S.A.-S. and A.A.-R.M.; investigation, S.A.-S. and A.A.-R.M.; resources, S.A.-S., A.A.-R.M.; curation, S.A.-S., Y.S. and B.B.; writing—original draft preparation, S.A.-S.; writing—review and editing, S.A.-S., Y.S., B.B.; visualization, S.A.-S.; B.B. and I.M, supervision; S.A.-S.; project administration, S.A.-S.; funding acquisition, S.A.-S. All authors have read and agreed to the published version of the manuscript.

## Institutional Review Board Statement

The study was authorized by the King Abdullah University Hospital’s (KAUH) institutional review board committee (20/51/2012). This study was carried out in compliance with the Helsinki Declaration.

## Informed Consent Statement

Clinical samples were obtained and donated from within stored clinical settings and therefore no data was stored regarding sample identification

## Data Availability Statement

Data is contained within the article or supplementary materials are also available upon request.

## Disclaimer

Statements contained within do not represent the views or opinions of authors and are accurate to the best of our knowledge based on scientific analysis and subject to local jurisdiction or advice from authorities or clinicians or references contained within.

