## Supplementary material for "A Genetic and Immunohistochemical Analysis of *H. Pylori* Phenotypes and p27 expression in Adenocarcinoma Patients in Jordan": Send to Brent-supplementary.pdf (from another version of this preprint: version 1, 2022-12-19)

*Supplementary Data*

### The Association between P27 Expression and *Helicobacter pylori* Infection in Gastric Adenocarcinoma Patients in Jordan

**Suhaila A. Al-Sheboul [1,*], Ahmad Abdul-Razzak Mohammad [1], Yasemin Shboul [1], Brent Brown [2], and Ismail I. Matalka [3]**

**Table S1**: Results of IHC staining of gastric cancer patients.

| | | Samples | Percent |
|---|---|---|---|
| IHC | Negative | 56 | 72.7% |
| | Positive | 21 | 27.3% |
| | Total | 77 | 100% |

**Table S2:** Distribution of p27 positive IHC results among various forms of gastric cancer.

| Type of Cancer | P27 Detection | | Total |
|---|---|---|---|
| | Negative | Positive | |
| Intestinal | 30 | 7 | 37 |
| | 81.1% | 18.9% | |
| Diffused | 26 | 14 | 40 |
| | 65.0% | 35.0% | |

**Table S3:** Distribution of the ureA gene sequence in patients with gastric cancer.

| | | Frequency | Percent |
|---|---|---|---|
| ***ureA* gene** | Negative | 35 | 45.5% |
| | Positive | 42 | 54.5% |
| | Total | 77 | 100.0% |

**Table S4:** Correlation between the IHC results and the PCR of the ureA gene sequence.

| **P27 expression** | ***H.pylori* Detection** | | **Total** |
|---|---|---|---|
| | **Negative** | **Positive** | |
| IHC Negative for p27 | 19 | 37 | 56 |
| | 33.9% | 66.1% | |
| IHC Positive for p27 | 16 | 5 | 21 |
| | 76.2% | 23.8% | |

**Table S5:** Distribution of cagA gene sequence of *H. pylori*-positive gastric cancer patients.

| | | **Frequency** | **Percent** |
|---|---|---|---|
| ***cagA* gene** | **Negative** | 18 | 23.4% |
| | **Positive** | 24 | 31.2% |
| | **Total** | 42 | 54.5% |
| [ | | | |

**Table S6:** Association between PCR of *cagA* gene sequence and IHC results

| **P27 expression** | ***cagA* detection** | | **Total** |
|---|---|---|---|
| | **Negative** | **Positive** | |
| IHC Negative for p27 | 14 | 23 | 37 |
| | 37.8% | 62.2% | |
| IHC Positive for p27 | 4 | 1 | 5 |
| | 80.0% | 20.0% | |

**Table S7:** Distribution of the vacAs1 gene sequence in gastric cancer patients with *H. pylori* infection**.**

| | | **Frequency** | **Percent** |
|---|---|---|---|
| ***vacAs1* gene** | **Negative** | 23 | 29.9% |
| | **Positive** | 19 | 24.7% |

| | | | |
|---|---|---|---|
| | Total | 42 | 54.5% |

**Table S8:** Correlation between PCR of *vacA*s1gene sequence and IHC results

| P27 expression | *vacA*s1 Detection | | Total |
|---|---|---|---|
| | Negative | Positive | |
| IHC Negative for p27 | 19 | 18 | 37 |
| | 51.4% | 48.6% | |
| IHC Positive for p27 | 4 | 1 | 5 |
| | 80.0% | 20.0% | |

**Table S9:** Distribution of *vacA*m1 gene sequence within gastric cancer patients positive for *H.pylori*

| | | Frequency | Percent |
|---|---|---|---|
| ***vacA*m1 gene** | **Negative** | 31 | 40.3% |
| | **Positive** | 11 | 14.3% |
| | **Total** | 42 | 54.5% |

**Table S10:** Distribution of positive *vacA*m1 gene in different types of gastric cancer.

| Type of Cancer | *vacA*m1 Detection | | Total |
|---|---|---|---|
| | Negative | Positive | |
| Intestinal | 16 | 4 | 20 |
| | 80.0% | 20.0% | |
| Diffused | 15 | 7 | 22 |
| | 68.2% | 31.8% | |

**Table S11:** Correlation between PCR of *vacA*m1 gene sequence and IHC results.

| P27 expression | *vacA*m1 Detection | Total |
|---|---|---|

| | **Negative** | **Positive** | |
|---|---|---|---|
| IHC Negative for p27 | 26 | 11 | 37 |
| | 70.3% | 29.7% | |
| IHC Positive for p27 | 5 | 0 | 5 |
| | 100.0% | .0% | |

**Table S12:** Distribution of *vacA*m2 gene sequence within gastric cancer patients positive for *H.pylori*.

| | | Frequency | Percent |
|---|---|---|---|
| ***vacA*m2 gene** | Negative | 31 | 40.3% |
| | Positive | 11 | 14.3% |
| | Total | 42 | 54.5% |
